# Place Cells in the Claustrum Remap Under NMDA Receptor Control

**DOI:** 10.1101/2021.04.21.440764

**Authors:** Emanuela Rizzello, Seán K. Martin, Jennifer Rouine, Charlotte K. Callaghan, Shane M. O’Mara

**Author notes:** **Corresponding authors**: Emanuela Rizzello, or Shane O’Mara, Trinity College Institute of Neuroscience, Trinity College Dublin, College Green, Ireland. **Conflict of interest**: The authors declare no competing financial interests.

## Abstract

Place cells are cells exhibiting location-dependent responses; they have mostly been studied in the hippocampus. Place cells have also been reported in the rat claustrum, an underexplored paracortical region with extensive corto-cortical connectivity. It has been hypothesised that claustral neuronal responses are anchored to cortical visual inputs. We show rat claustral place cells remap when visual inputs are eliminated from the environment and that this remapping is NMDA-receptor-dependent. Eliminating visual input enhances delta-band oscillatory activity in the claustrum, without affecting simultaneously-recorded visual cortical activity. We conclude that, like the hippocampus, claustral place field remapping might be mediated by NMDA receptor activity, and is modulated by visual cortical inputs. (106 words)

## Introduction

In mammals, several brain regions (including the hippocampus, anterior thalamus, and claustrum) possess cells the firing of which is localized to ‘place fields’ which code the animal’s position in an environment (see Grieves and Jeffery, 2017; O’Mara and Aggleton, 2019 for review). Place cells have been mostly investigated in the hippocampus, and there are few reports of extra-hippocampal place cells (Grieves and Jeffery, 2017). Jankowski and O’Mara (2015) reported spatially-coding neurons responding to position, boundaries, object location and head direction, across light and dark conditions in the claustrum; moreover, claustral place cells also responded to the rotation of visual distal cues in the environment.

Compared to the hippocampus, the claustrum is an under-investigated paracortical region; it is adjacent the orbitofrontal cortex (anteriorly), insular cortex (laterally), and the caudate nucleus of the striatum (medially). The claustrum is comprised of a thin sheet of neurons, possessing a complex dorsoventral and rostrocaudal topography largely spanning the rostral half of the telencephalon (Dillingham *et al*., 2017). There are extensive connections between claustrum and visual cortex (Pearson *et al*., 1982), including dense ipsilateral inputs from the secondary visual cortex (Miller and Vogt, 1984). We therefore reasoned claustral place field remapping might be affected by visual inputs. We manipulated visual inputs during freely-moving navigation by foraging rats to explore potential mechanisms underpinning claustral place fields. We also measured oscillatory activity in the claustrum and visual cortex during light/dark manipulation. Place fields remained present but rapidly remapped by about 50^0^ when visual inputs were eliminated by turning off the room lights. Previous experiments have shown place field remapping in the hippocampus is under NMDA receptor control (Kentros *et al*., 1998). We therefore modulated NMDA receptor activity via systemic D-serine administration (an NMDA-receptor agonist that readily passes the blood-brain-barrier), finding the expected 50^0^ remapping of claustral place cells when ambient visual inputs are eliminated was absent. We also found that darkness exposure alters claustral oscillations in the delta-band, while preserving visual cortical activity. Further, the presence of a distal beacon induced changes in the visual cortex and claustrum which are absent during the light/dark protocol. We conclude that, like the hippocampus, claustral place field remapping might be mediated by NMDA receptor activity, and is modulated by visual cortical inputs. (371 words)

## Results

After post-mortem histological verification, 52 well-isolated units recorded in 13 rats were assigned to the claustrum. Based on their electrophysiological and spatial properties, 43 units were classified as place cells (82.69%), and 9 as bursting cells without spatially-related firing (36.54%; Table 1)

**Table 1.** Electrophysiological classification of claustral units. Summary statistics for mean spike width, mean spike amplitude, mean inter-spike interval (ISI), mean spike frequency and spatial Information content (Skaggs) (mean ± SEM) for all claustral place and bursting cells recorded in the claustrum.

|  | Place cells | Bursting cells |
| --- | --- | --- |
| Width (μs) | 249.20 ± 9.01 | 253.00 ± 84.33 |
| Amplitude (μV) | 134.89 ± 19.76 | 126.03 ± 42.01 |
| ISI (mS) | 3869.03 ± 968.24 | 1514.05 ± 618.10 |
| Spike freq. (Hz) | 0.34 ± 0.09 | 3.06 ± 1.02 |
| Spatial Skaggs | 2.54 ± 0.29 | 1.48 ± 0.49 |

We performed recordings (see Methods) in which rats navigated in a square arena while foraging for food pellets. Additionally, we investigated the effects of light/dark transitions to explore how claustral places are modulated by changes in ambient visual inputs (i.e. light or dark during the recording session; Brotons-Mas *et al*., 2010). In the light condition, the arena was indirectly lit by four symmetrically-positioned spotlights. The animal navigated in a square arena for 30 minutes; visual inputs changed every 10 minutes from light (L1) to dark (D) and back to light (L2). At the end of each session the arena was cleaned to remove odours. The next day (Day 2) rats received a subcutaneous injection of D-SER (n = 6 rats) thirty minutes before recording. Recordings were repeated at the same time as Day 1 following the same procedure (L1 _D-SER_, D _D-SER_, L2 _D-SER_). This protocol was used to measure the place field stability/remapping between different conditions, and their modulation by NMDA receptor activity. The D-SER effect was also assessed via oscillatory power (total power, subgroup, n = 3), before and after drug administration.

### Claustral place cells remap in darkness

We found place fields were stable between the two sessions in light (before and after dark exposure). When visual inputs were eliminated from the environment, place fields remapped their position consistently and significantly. The measure of the distance between the centre of mass (COM) of each place field in the three different conditions (L1, D and L2) revealed a shift of the place field from light to dark (L1-D). The place field returned to a similar position in the first light exposure during the last recording in light (L2) (ΔL1-D (cm) = 24.97 ± 5.50, ΔL1-L2 (cm) 18.73 ± 5.20, ΔD-L2 (cm) = 16.60 ± 4.29, mean ± SEM, Fig.1d). A consistent angle was formed by the three place fields (L1, D and L2) in the arena (Fig. 1b). We used the centre of mass of the place fields (x and y coordinates) to measure potential differences between the three conditions (see Methods for details). Notably, we found claustral place cells consistently remapped by about 50 degrees in darkness (L1DL2 angle (degrees): 47.84 ± 6.82, n=19, mean ± SEM, Fig. 1c), compared to light conditions.

**Fig. 1.**
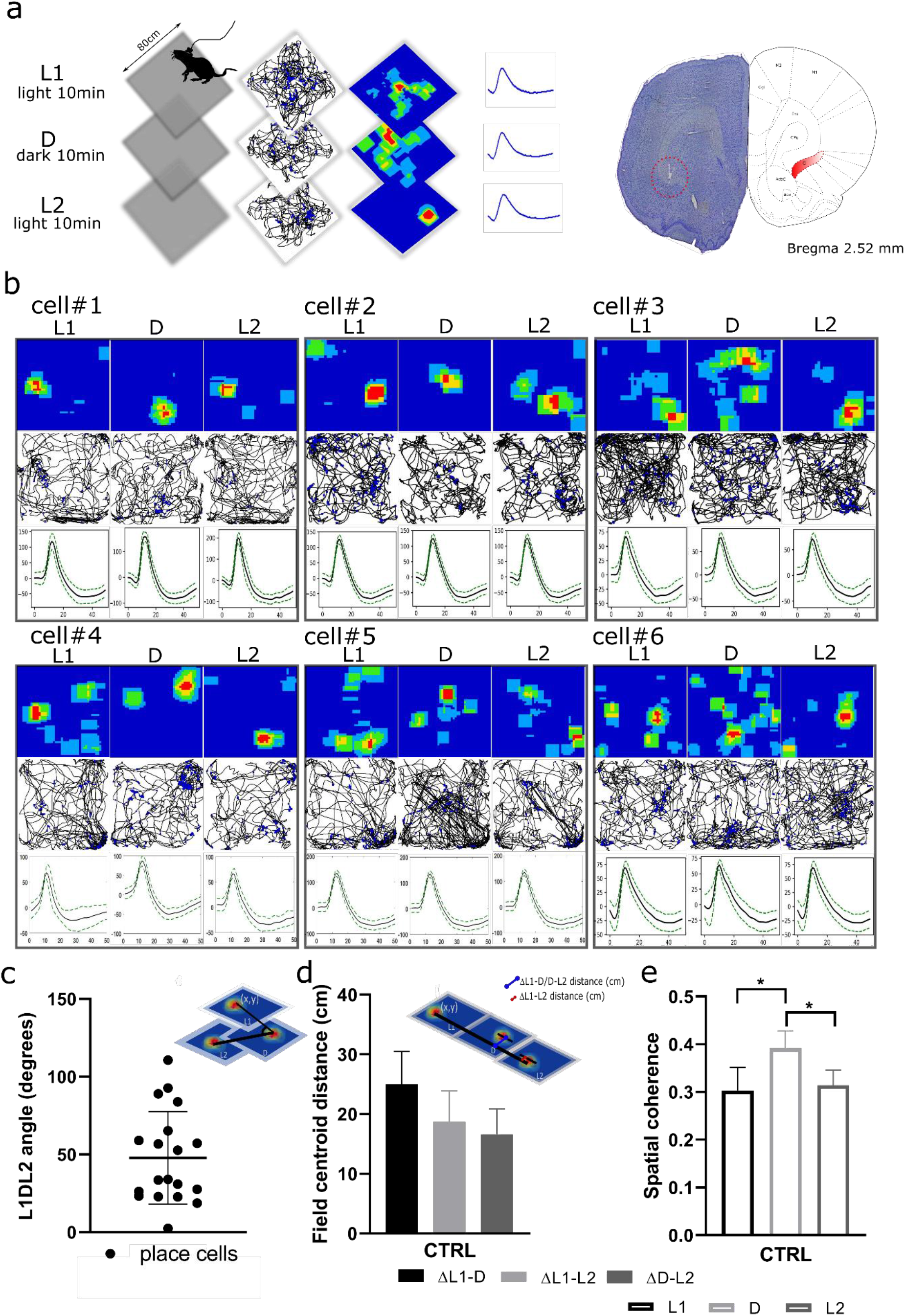
Claustral place cells remap in darkness. (a) On Day 1, claustral place cells were recorded over consecutive sessions (10 minutes each), during which animals navigated in the square arena (80×80 cm) with light manipulation (light (L1)/dark (D)/light (L2)); the figure shows an example of the animal path recorded during navigation with superimposed firing activity of the unit (black line with blue dots), the firing rate map (blue square) and a single unit waveform corresponding to a claustral place cell. On completion of the recording studies positions of recorded cells were estimated by histological analyses (the red circle shows tetrodes in the claustrum). (b) Examples of six claustral place cells recorded over consecutive sessions with changes in the environmental conditions; when the light was eliminated from the environment, a shift of the claustral place field occurred with a L1DL2 angle (c) of about 50 degrees. (d) The measure of the distance between the center of mass (COM) of each place field in three different conditions (L1, D and L2) revealed a shift of the place field in the dark. (e) Spatial coherence was statistically different between light and dark conditions, and there were no differences between the first and the last light exposure.

Overall, our results show that when visual inputs are removed the claustral place map moves to a new constant location in the arena in darkness.

### NMDA receptor modulation sharply changes claustral place cell remapping

We tested whether the modulation of NMDA receptor activity decreased remapping in the dark. Firstly, thirty minutes before the recording session of Day 2, we subcutaneously injected the NMDA receptor co-agonist D-SER (see Methods). The experimental procedure of Day 2 (day in which rats received the drug) was the same as Day 1 (L1/D/L2), except the drug pre-injection administration during a time window where the drug shows its highest peak of activity (Fig. 2a). We found again the claustral place cells consistently remapped by about 50 degrees in darkness, but that the angle was significantly increased following D-SER injection in each cell recorded (L1DL2 angle _CTRL_ (degrees) = 47.57 ± 11.39 vs L1DL2 angle _D SER_ = 104.13 ± 17.82, p= 0.003, n=9, mean ± SEM, two-tailed *t* test for paired two samples for means, Fig. 2c). The increase in the angle indicates that the three place fields were formed at a similar location. In D-SER treated rats the measure of the difference in terms of distance between the place field CoM in L1, D and L2 revealed the three place fields at a similar location (ΔL1-D (cm) = 13.82 ± 4.84, ΔL1-L2 (cm) 20.30 ± 4.81, ΔD-L2 (cm) = 14.74 ± 3.47, mean ± SEM, Fig. 2d).

**Fig. 2.**
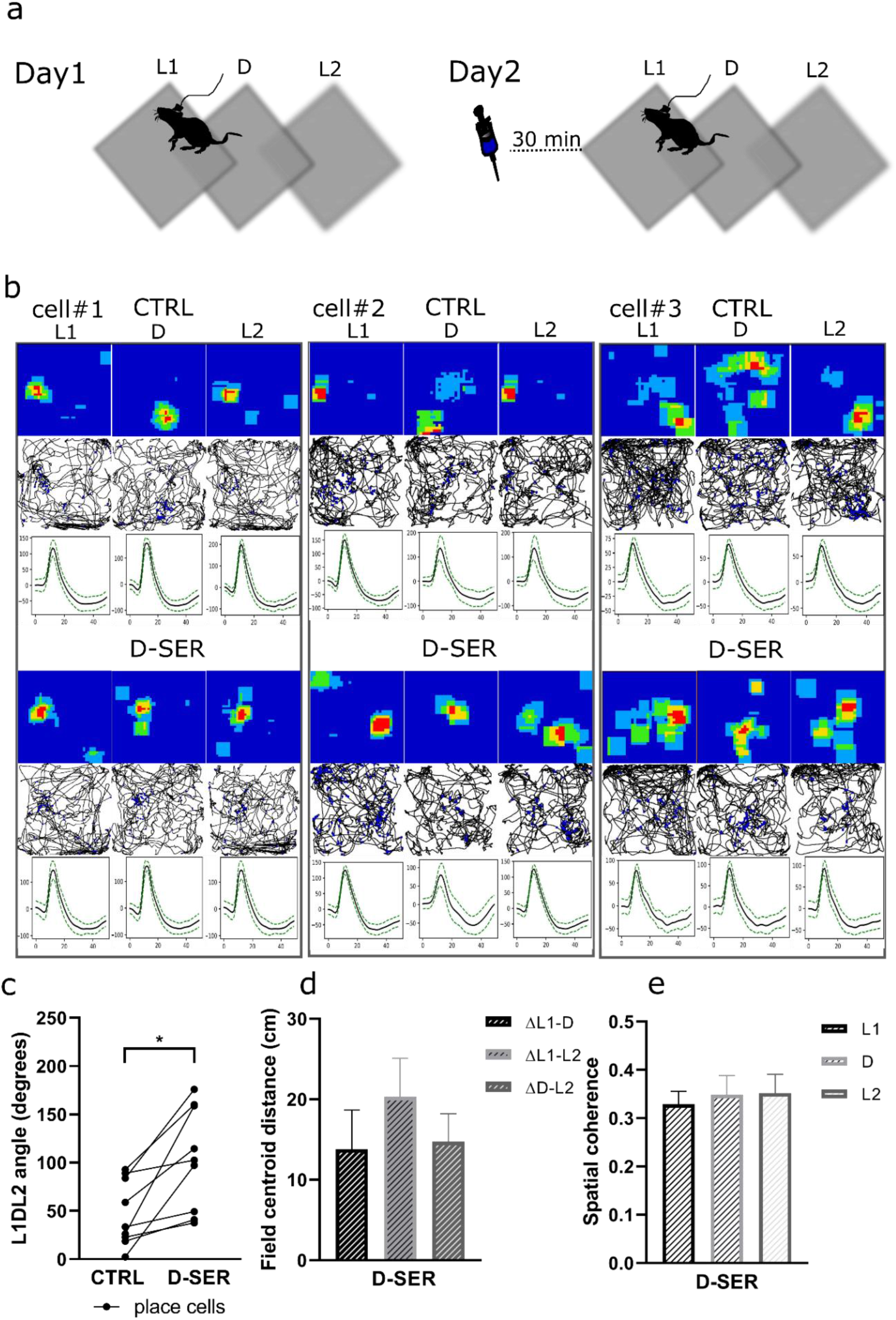
D-SER reduces claustral place cells remapping. (a) Day 2 replicated Day 1 but included subcutaneous administration of D-SER (10mg/Kg). (b) Example of three claustral place cells in ctrl and following D-SER administration; place cells remapping in dark was reduced following D-SER administration. (c) L1DL2 angle and (d) distance increased following D-SER administration. (e) Spatial coherence was not statistically different between light and dark following D-SER administration.

We measured spatial (Skaggs) information content in controls, and following drug administration, in each condition. Consistent with Jankowski and O’Mara (2015), we found that the spatial information content was significantly higher in the dark (D _CTRL_ = 3.09 ± 0.32, mean ± SEM), compared to the preceding recording in light (L1 _CTRL_ = 2.54 ± 0.29, p =0.012, mean ± SEM, two-tailed t test paired). Contrarily, we found a difference in spatial information content between the recordings in the dark (D _CTRL_ = 3.09± 0.32, mean ± SEM) and the following recordings in the light (L2 _CTRL_ = 2.36 ± 0.34, p=0.006, mean ± SEM, two-tailed t test paired). There were no significant differences in spatial information content between the first and the last recording in light (L1 _CTRL_ vs L2 _CTRL_ p = 0.06, two-tailed t test, Fig. 1e). Interestingly, there were no significant differences in spatial information content between L1, D and L2 following D-SER administration (L1 _D-SER_ = 3.17 ± 0.43 vs D _D-SER_ = 3.14 ± 0.29, p = 0.93; D _D-SER_ = 3.14 ± 0.29 vs L2 _D-SER_ = 3.12 ± 0.31, p = 0.92; L1 _D-SER_ = 3.17 ± 0.43 vs L2 _D-SER_ = 3.12 ± 0.31, p = 0.92, n = 9, mean ± SEM, two-tailed t test paired, Fig. 2e).

Place field remapping was assessed by measuring spatial coherence across conditions; this revealed a significant difference between the first light and dark (L1 _CTRL_ = 0.30 ± 0.05 vs D _CTRL_ = 0.39 ± 0.04, p = 0.05, Fig. 1e) and between dark and the last light (D _CTRL_ = 0.39 ± 0.04 vs L2 _CTRL_ = 0.31 ± 0.03, p = 0.01). There was no difference in spatial coherence between first and last light exposure (L1 _CTRL_ = 0.30 ± 0.04 vs L2 _CTRL_ = 0.31 ± 0.03, p = 0.73, mean ± SEM, two-tailed t test paired, Fig. 1e) suggesting place field remapping just occurred in darkness. The difference in the spatial coherence disappeared following D-SER injection between the first light and dark (L1 _D-SER_ = 0.32 ± 0.02 vs D _D-SER_ = 0.35 ± 0.04, p = 0.52, n = 9, mean ± SEM, two-tailed t test paired, Fig. 2e) and between dark and the last light (D _D-SER_ = 0.35 ± 0.04 vs L2 _D-SER_ = 0.35 ± 0.04, p = 0.94, n = 9, mean ± SEM, two-tailed t test paired). D-SER administration had no effect in the spatial coherence similarity between the first and last light (L1 _D-SER_ = 0.32 ± 0.02 vs L2 _D-SER_ = 0.35 ± 0.04, p = 0.42, n = 9, mean ± SEM, two-tailed t test paired). These results suggest that D-SER decreased the place field remapping in darkness.

### Single unit firing properties are modulated by NMDA receptor activation

We next investigated whether the drug effect on place field remapping in darkness underlies the firing changes of single place cells in the claustrum. To this end, we measured both the action potential and firing properties in controls and following drug administration. Previous studies demonstrated that place cell firing properties change between light and dark in the hippocampus. However, we observed no statistically significant differences in the control recordings between L1/D/L2 transition for spike amplitude (L1 _CTRL_ (µV) = 134.90± 19.76 vs D _CTRL_ = 153.38 ± 21.73, p = 0.22, n =9; D _CTRL_ = 153.38 ± 21.73 vs L2 _CTRL_ = 122.51 ± 12.46, p = 0.06, n=9; L1 _CTRL_ = 134.90± 19.76 vs L2 _CTRL_ = 122.51 ± 12.46, p = 0.31, n=9, mean ± SEM, two-tailed t test paired), and spike width (L1 _CTRL_ (µs) = 249.20 ± 9.01 vs D _CTRL_ = 245.48 ± 19.04, p = 0.82, n =9; D _CTRL_ = 245.48 ± 19.04 vs L2 _CTRL_ = 257.11 ± 14.55, p = 0.34, n=9; L1 _CTRL_ = 249.20 ± 9.01 vs L2 _CTRL_ = 257.11 ± 14.55, p = 0.46, n=9, mean ± SEM, two-tailed t test paired). Interestingly, spike frequency was significantly lower in the dark, compared to the preceding recording in light (L1 _CTRL_ (Hz) = 0.35 ± 0.09 vs D _CTRL_ = 0.23 ± 0.09, p= 0.04, n = 9, mean ± SEM, two-tailed t test paired, Fig.3a), as well as the following recording in light (D _CTRL_ (Hz) = 0.23 ± 0.09 vs L2 _CTRL_ = 0.39 ± 0.08, p= 0.02, n= 9, mean ± SEM, two-tailed t test paired). There were no significant differences in the spike frequency between the first and final recording in light (L1 _CTRL_ (Hz) = 0.35 ± 0.09 vs L2 _CTRL_ = 0.39 ± 0.08, p = 0.08, n = 9, mean ± SEM, two-tailed t test paired).

**Fig. 3.**
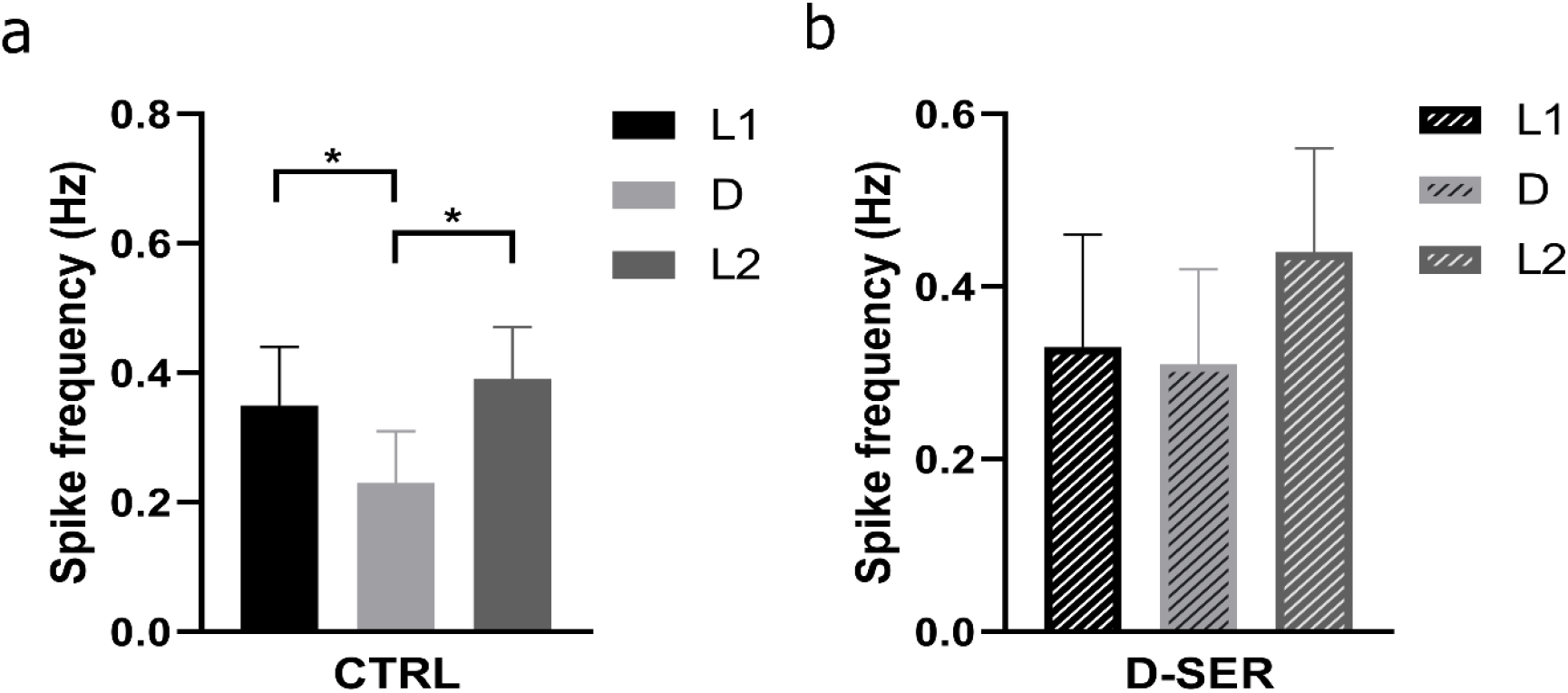
Claustral place field remapping underlies a change of excitability. (a) The exposure to darkness significantly decreased claustral place cells excitability. (b) D-SER removed differences in the spike frequency between light and dark.

These findings raise the question: is the decrease in remapping by D-SER caused by a change of excitability? To address this issue, we analysed spike frequency following D-SER injection. Notably, D-SER removed the statistical difference between L1/D/L2 in spike frequency (L1 _D-SER_ (Hz) = 0.33 ± 0.13 vs D _D-SER_ = 0.31 ± 0.11, p = 0.75, n = 9, mean ± SEM, two-tailed t test paired, Fig.3b; D _D-SER_ = 0.31 ± 0.11 vs L2 _D-SER_ = 0.44 ± 0.12, p = 0.13, n= 9, mean ± SEM, two-tailed t test paired; L1 _D-SER_ = 0.33 ± 0.13 vs L2 _D-SER_ = 0.44 ± 0.12, p= 0.27, n= 9, mean ± SEM, two-tailed t test paired, Fig.3b), suggesting the decrease in place field remapping following D-SER injection might result from an increase in place cell excitability in darkness.

### Visual manipulations do not interfere with bursting cell activity

We next investigated whether the influence of the visual input elimination was extended to other cells observed in the claustrum. We found that the 36.54% of the cell population in the claustrum were bursting cells without spatially-related firing. Following their characterization (Table 1), we analysed spike properties and firing rate of bursting cells over the L1/D/L2 sequence. There were no difference in the spike width over the three conditions (L1 (µs) = 253.00 ± 84.33 vs D = 254.13 ± 84.71, p = 0.78, n = 9; D = 254.13 ± 84.71 vs L2 = 251.94 ± 83.98, p = 0.58, n = 9; L1 = 253.00 ± 84.33 vs L2 = 251.94 ± 83.98, p = 0.79, n = 9, mean ± SEM, two-tailed t test paired). Similarly, no statistically significant differences were observed in the spike amplitude over the three conditions (L1 (µV) = 126.04 ± 42.01 vs D = 120.31 ± 40.10, p = 0.23, n = 9; D = 120.31 ± 40.10 vs L2 = 124.81 ± 41.60, p = 0.28, n = 9; L1 = 126.04 ± 42.01 vs L2 = 124.81 ± 41.60, p = 0.69, n = 9, mean ± SEM, two-tailed t test paired). The light/dark contrasts had no effect on the spike frequency of bursting cells (L1 (Hz) = 3.06 ± 1.02 vs D = 3.01 ± 1.00, p = 0.93, n = 9; D = 3.01 ± 1.00 vs L2 = 3.03 ± 1.00, p = 0.97, n = 9; L1 = 3.06 ± 1.02 vs L2 = 3.03 ± 1.00, p = 0.91, n = 9, mean ± SEM, two-tailed t test paired). Finally, we measured the total number of burst events for each cell, and observed that visual stimuli did not affect bursting activity (L1 = 60.33 ± 20.11 vs D = 45.22 ± 15.07, p = 0.45, n = 9; D = 45.22 ± 15.07 vs L2 = 65.38 ± 23.11, p = 0.29, n = 9; L1 = 60.33 ± 20.11 vs L2 = 65.38 ± 23.11, p = 0.42, n = 9, mean ± SEM, two-tailed t test paired). Thus, the manipulation of visual stimuli in the environment does not affect spike and firing properties of claustral bursting cells.

### Locomotor activity is not influenced by D-SER administration

Place cell firing rate is positively correlated with the animals’ running speed. To determine whether the firing rate modulation by D-SER correlates with changes in running speed, we measured average running speeds before and after drug administration (L1 _CTRL_ vs L1 _D-SER_). Our data suggest that D-SER did not influence the animal behaviour during navigation (L1 _CTRL_ (cm/sec) = 10.05 ± 4.10 vs L1 _D-SER_ = 9.47 ± 3.87, n rats = 6, p = 0.71, n.s. mean ± SEM, two-tailed t test paired), suggesting D-SER did not affect running speed.

### Claustral theta oscillations are not affected by changes to visual inputs

To determine how theta oscillations are modulated by dark exposure, we measured the relative power of theta oscillations over the L1/D/L2 sequence, and compared how NMDA receptor activation by the co-agonist D-SER influences the claustral local circuit and the neuronal response to light/dark transition. Specifically, we compared theta oscillations for control (Day 1), and following D-SER administration (Day 2), and across the two days of recording (Day1 vs Day2). Visual input manipulations had no effect on oscillations (L1 _CTRL_ = 0.36 ± 0.02 vs D _CTRL_ = 0.37 ± 0.03, p = 0.72, n = 9; D _CTRL_ = 0.37 ± 0.03 vs L2 _CTRL_ = 0.36 ± 0.02, p = 0.63, n = 9; L1 _CTRL_ = 0.36 ± 0.02 vs L2 _CTRL_ = 0.36 ± 0.02, p = 0.87, n = 9, mean ± SEM, two-tailed t test paired, Fig. 4a). Likewise, D-SER administration did not affect the relative power of theta oscillations over the L1/D/L2 sequence (L1 _D-SER_ = 0.34 ± 0.02 vs D _D-SER_ = 0.33 ± 0.02, p = 0.53, n = 9; D _D-SER_ = 0.33 ± 0.02 vs L2 _D-SER_ = 0.32 ± 0.02, p = 0.27, n = 9; L1 _D-SER_ = 0.34 ± 0.02 vs L2 _D-SER_ = 0.32 ± 0.02, p = 0.21, n = 9, mean ± SEM, two-tailed t test paired, Fig. 4a). Although a visible theta oscillation reduction was found by comparing each subset of visual stimuli input before and after drug administration, the overall difference was not statistically significant (L1 _CTRL_ = 0.36 ± 0.02 vs L1 _D-SER_ = 0.34 ± 0.02, p = 0.074, n = 9; D _CTRL_ = 0.37 ± 0.03 vs D _D-SER_ = 0.33 ± 0.02, p = 0.074, n = 9; L2 _CTRL_ = 0.36 ± 0.02 vs L2 _D-SER_ = 0.32 ± 0.02, p = 0.055, n = 9, mean ± SEM, Wilcoxon matched-pairs signed rank test, Fig. 4a, b, c). The calculation of a p value in the Wilcoxon matched-pairs signed rank test is inaccurate where the number of test samples is less than 10. Since N is smaller than 10 in this study, we compared the signed rank to the critical value for the sample size at the 5% significance level. For example, in the Wilcoxon test for differences in theta oscillations between L2 _CTRL_ and L2 _D-SER_, the signed rank was 33, which was below the critical value of 35 necessary for statistical significance. These results suggest that the theta oscillations do not correlate with claustral place cell remapping in darkness.

**Fig. 4.**
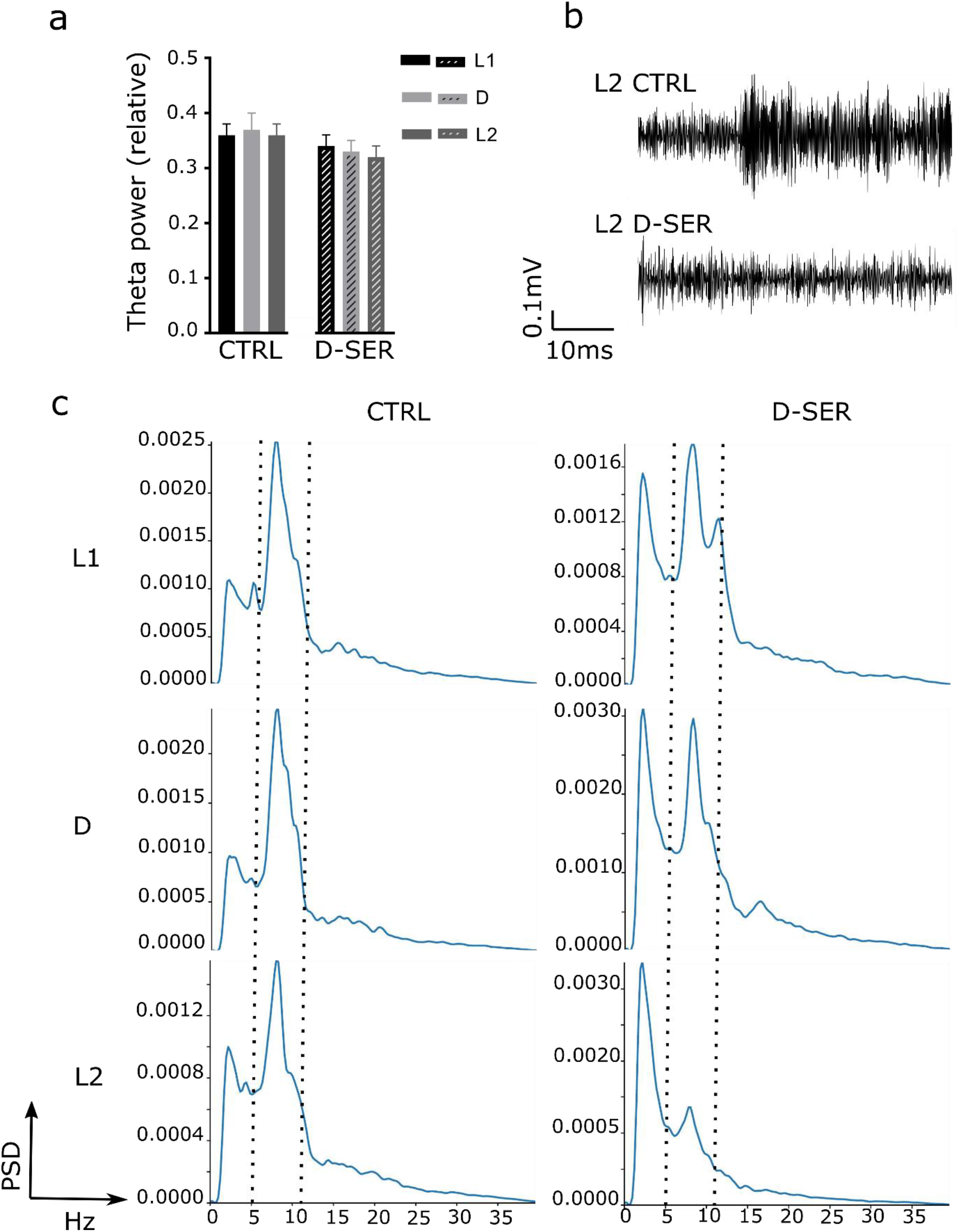
Theta oscillations following light and NMDAR manipulation. (a) Theta power did not change following visual manipulation; D-SER administration did not affect theta power oscillations along the L1/D/L2 sequence. (b) Example of LFP traces in the theta range (50ms selected from 10 min of LFP recording) in control and following D-SER administration during recordings in L2. (c) Periodogram in L1/D/L2 in control, and following D-SER administration (theta range is highlighted by the black dashed).

Then, to determine how effectively the 10mg/Kg dose D-SER influences the local claustral network, we examined total power following drug administration. The administration of D-SER significantly increased the total power in the claustrum (total power saline (mV^2^) = 0.0047 ± 0.000269 vs total power D-SER (mV^2^) = 0.0053 ± 0.000339, p = 0.000001, n channels = 24, n rats = 3, mean ± SEM, two-tailed t test paired, Fig.5). These results show that the dose of D-SER used for place cell recordings effectively activated NMDAR receptors, increasing claustral neuronal excitability.

**Fig. 5.**
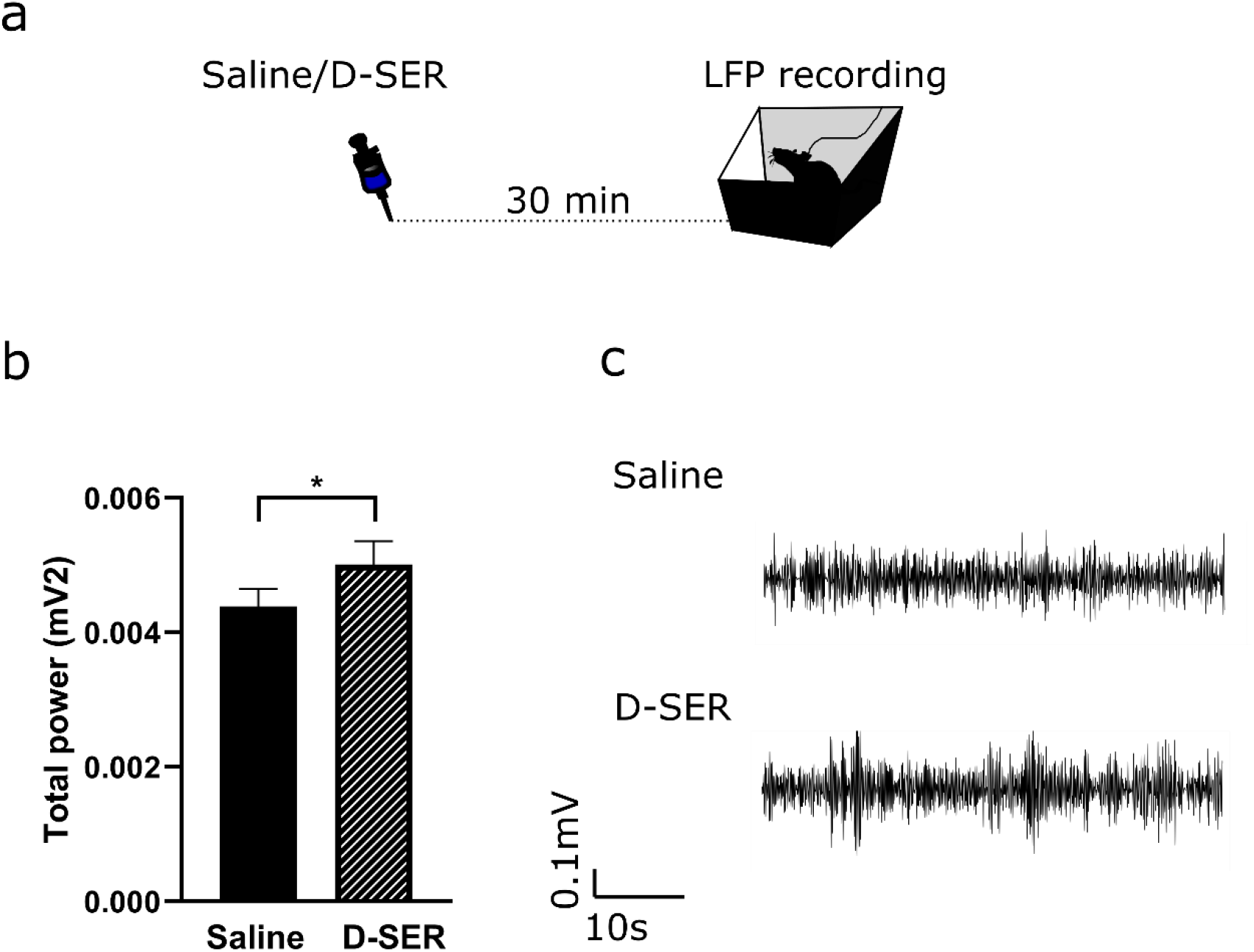
Total power increases following D-SER administration. (a) LFPs were recorded following a subcutaneous injection of normal saline (Day1) or D-SER (Day2) in a box (24h x 38w x 38l cm). (b) D-SER significantly increases the total power in the claustrum. (c) Representative traces (50 ms segments extracted from 10 min recording) of the total power in saline (top) and following D-SER administration (bottom).

### Claustral delta oscillations change from light to dark

The constant, reliable angle formed by remapping claustral place fields in darkness led us to hypothesize that remapping arose from a loss of inputs from visual cortices to the claustrum, and therefore a loss of visual calibration to support place field stability. The claustrum receives dense ipsilateral inputs from visual cortices, and projects back to these areas. We therefore recorded oscillatory activity in secondary visual cortex and claustrum during navigation in light and dark.

The first question was to investigate whether the protocol that showed us the remapping caused changes in both visual and claustral oscillatory activity. Following tetrode implantation in these two brain areas, animals navigated in the square arena moving from light (L1) to dark (D) (Fig. 6a) while the local field potential was simultaneously recorded. The total power of raw LFP oscillations in the V2L cortex as well as the relative powers in V2L of δ, θ and γ oscillations were no different between light and dark (L1_tot_ (mV^2^) = 0.008 ± 0.001 vs D_tot_ (mV^2^) = 0.008 ± 0.001, increased by 2.95%; L1_δ_ (mV^2^) = 0.0012 ± 0.0002 vs D_δ_ (mV^2^) = 0.0012 ± 0.0002, decreased by 4.66%; L1_θ_ (mV^2^) = 0.0044 ± 0.0008 vs D_θ_ (mV^2^) = 0.0047 ± 0.0010, increased by 7.90%; L1_γ_ (mV^2^) = 0.00055 ± 0.00001 vs D_γ_ (mV^2^) = 0.00057 ± 0.00003, increased by 4.26%, n samples = 4, Fig.6b). The total power of raw LFP oscillations in the claustrum was no different between light and dark (L1_tot_ (mV^2^) = 0.006 ± 0.001 vs D_tot_ (mV^2^) = 0.006 ± 0.001, decreased by 1.72%, n samples = 4, Fig. 6c). The relative powers in claustrum of θ and γ oscillations were no different between light and dark (L1_θ_ (mV^2^) = 0.0017 ± 0.0008 vs D_θ_ (mV^2^) = 0.002 ± 0.001, increased by 9.91%; L1_γ_ (mV^2^) = 0.00056 ± 0.00001 vs D_γ_ (mV^2^) = 0.00059 ± 0.00003, increased by 4.15%, n samples = 4, Fig. 6c). Interestingly, the relative power in claustrum of δ oscillations were consistently decreased in dark compared to that in light (L1_δ_ (mV^2^) = 0.0017 ± 0.0003 vs D_δ_ (mV^2^) = 0.0014 ± 0.0002, decreased by 17.54%, n samples = 4, Fig. 6c). Elimination of the light from the environment does not affect the secondary visual cortical activity, but triggers substantial changes in the delta-band oscillations of the claustrum. Claustral place cell remapping might be anchored to inputs unrelated to secondary visual cortical activity.

**Fig. 6.**
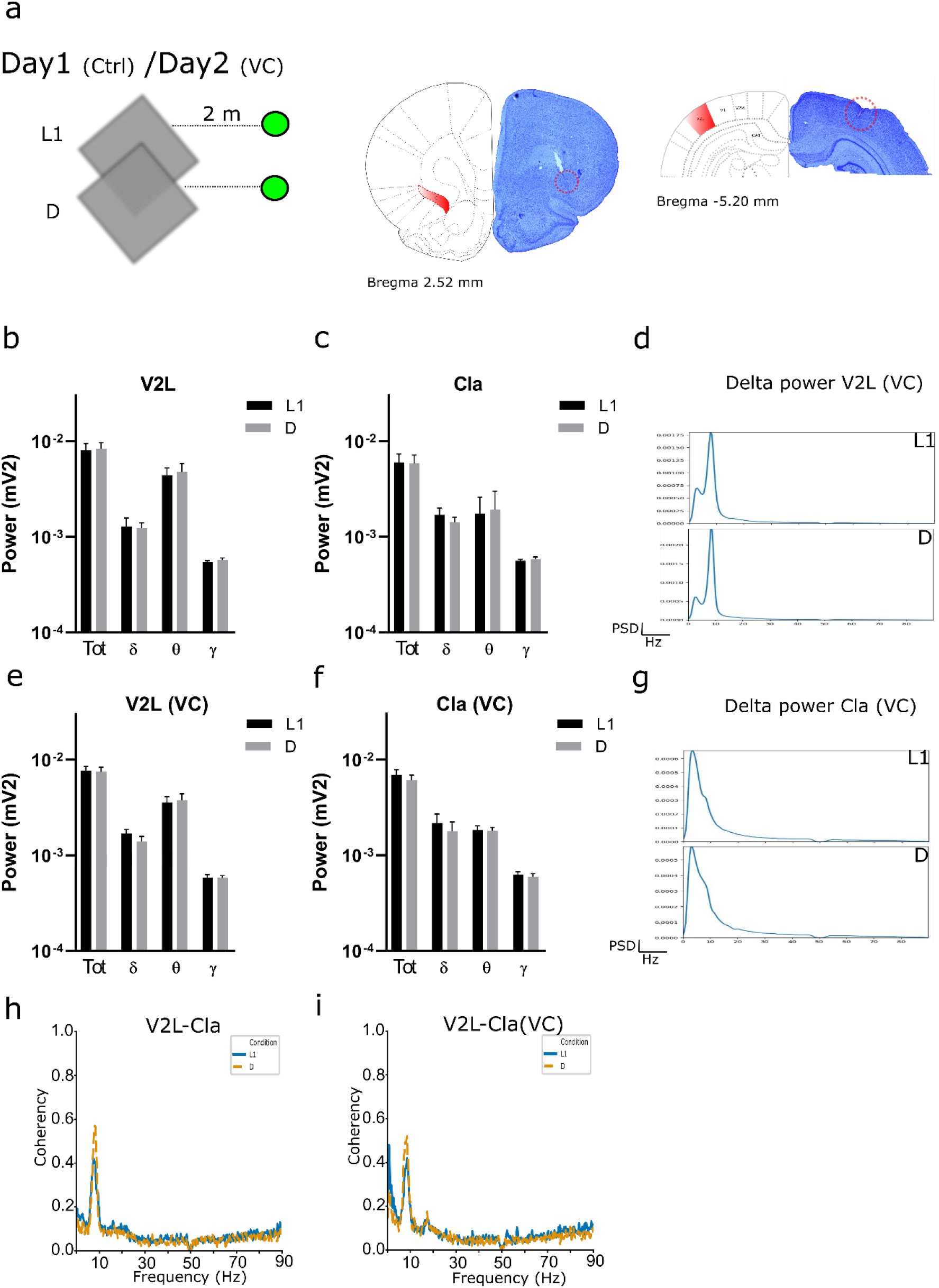
The distal, salient beacon modulates oscillations in claustrum and visual cortex. (a) The Day 1 animals navigated in a square arena moving from light (L1) to dark (D); the Day 2, the study of the Day 1 was repeated in the presence of a distal beacon (VC) in the environment; tetrodes and electrodes positions were estimated with histological analyses (the red circle shows tetrodes in the claustrum (left) and electrodes in V2L (right). Total and relative powers in V2L from L1 to D in the absence (b) or presence (e) of a distal beacon. Total and relative powers in claustrum from L1 to D in the absence (c) or presence (f) of a distal beacon. Periodogram in L1/D of the delta power from V2L (d) or claustrum (g) in the presence of a distal beacon. Coherency between V2L and claustrum in absence (h) or presence (i) of a distal beacon.

### Delta-band oscillations are highlighted by a distal beacon in the environment

The functional significance of delta oscillations in the brain has been linked to cognitive processes related to attention and detection of motivationally-salient stimuli in the environment (i.e. objects in the environment that attract attention). Claustral remapping was observed when the light was eliminated from the environment, and the concomitant reduction in the delta-band oscillations, might be a result of insufficient attention to distal objects or beacons in the environment used as anchors during navigation in light. To test this hypothesis, the previous protocol (light/dark) was repeated in the presence of a distal, salient beacon (flashing green light, Fig. 6a) in light and dark to attract the animal’s attention during navigation. The total power of raw LFP oscillations in the V2L cortex, as well as the relative powers in V2L of θ and γ oscillations were not different between light and dark in the presence of the visual distal cue (L1_tot_ (mV^2^) = 0.0077 ± 0.0008 vs D_tot_ (mV^2^) = 0.0074 ± 0.0009, decreased by 3.00%; L1_θ_ (mV^2^) = 0.0036 ± 0.0005 vs D_θ_ (mV^2^) = 0.0038 ± 0.0006, increased by 6.00%; L1_γ_ (mV^2^) = 0.00058 ± 0.00005 vs D_γ_ (mV^2^) = 0.0006 ± 0.00003, n samples = 4, Fig. 6e). Interestingly, the relative power in V2L of δ oscillations was consistently decreased in dark compared to that in light in the presence of the visual distal cue (L1_δ_ (mV^2^) = 0.0017 ± 0.0002 vs D_δ_ (mV^2^) = 0.0014 ± 0.0002, decreased by 19.05%, n samples = 4, Fig. 6d).

Correspondingly, the relative power in claustrum of δ oscillations was consistently decreased in dark, compared to that in the light in the presence of the visual distal cue (L1_δ_ (mV^2^) = 0.0022 ± 0.0005 vs D_δ_ (mV^2^) = 0.0012 ± 0.0004, decreased by 19.52%, n samples = 4, Fig. 6g). The total power of raw LFP oscillations in the claustrum as well as the relative powers of θ and γ oscillations was no different between light and dark in the presence of the distal beacon (L1_tot_ (mV^2^) = 0.006916 ± 0.00093401 vs D_tot_ (mV^2^) = 0.00609375 ± 0.00078946, decreased by 12%; L1_θ_ (mV^2^) = 0.0018325 ± 0.00020288 vs D_θ_ (mV^2^) = 0.0018053 ± 0.00015373, decreased by 1.50%; L1_γ_ (mV^2^) = 0.00062725 ± 0.000043175 vs D_γ_ (mV^2^) = 0.000595 ± 0.00004865, decreased by 5.28%, n samples = 4, Fig. 6f). These results confirmed our hypothesis that the presence of a distal, salient beacon in the environment might trigger changes in delta-band oscillatory activity, and this process might underlie the remapping phenomenon observed in the claustral place cells.

### Delta and Theta coherency is modulated by the distal, salient beacon

It has been suggested that coherent oscillations of neuronal activity might mediate interactions between different brain areas. We analyzed coherence between the secondary visual cortex and claustrum during navigation in light and dark for frequencies between 0 and 90 Hz. Coherence between these two brain areas was higher in the δ and θ bands in the presence and absence of the distal beacon (Coherence V2L-Cla L1δ = 0.2955; Dδ = 0.1824; L1θ = 0.4182; Dθ = 0.5195; L1γ = 0.1309; Dγ = 0.1054; Coherence V2L-Cla (VC) L1δ = 0.1700; Dδ = 0.1324; L1θ = 0.4225; Dθ = 0.5716; L1γ = 0.1318; Dγ = 0.1023, Fig. 6h and i). Notably, coherence between secondary visual cortex and claustrum was higher when the distal visual cue was introduced in the environment (Fig. 6i). These results confirm a possible communication between these two brain areas while revealing a remarkable contribution of δ and θ oscillations. The presence of the distal beacon in the environment does not alter the nature of this communication.

## Discussion

Here, we explored how claustral place cells remap when visual inputs are eliminated from the environment, finding they remap by a constant angle; this remapping is likely to be NMDA-receptor-dependent. Furthermore, eliminating visual inputs enhances delta-band oscillatory activity in the claustrum without affecting simultaneously-recorded visual cortical activity. We conclude that, like the hippocampus, claustral place field remapping might be mediated by NMDA receptor activity, and is modulated by visual cortical inputs. We also investigated claustral unit firing changes to explore the mechanism underlying claustral place field remapping in darkness. We established there was an absence of rate remapping between the two light conditions (i.e., before and following the dark condition). Intriguingly, there was a reduction in firing rates when visual inputs were eliminated from the environment, accompanied by place remapping in the dark.

Several lines of evidence suggest spatial maps (including their stability and expression) are influenced by plasticity-dependent mechanisms. NMDA receptor–dependent synaptic plasticity is necessary for establishing stable neuronal representations in the hippocampus (Kentros *et al*., 1998; Shapiro, 2001). Indeed, Shapiro (2001) concluded NMDA receptor antagonists prevent the establishment of stable place fields in the hippocampus. Furthermore, Kentros et al. (1998) observed that blocking NMDA receptors abolished the long-term stability of place field remapping in the hippocampus. Thus, to explore the mechanisms underlying claustral place field remapping in darkness, we administered the NMDA receptor co-agonist D-SER before performing the light/dark/light recording. Iin addition to glutamate, NMDA receptor activation needs the binding of a co-agonist at the GluN1 subunit (Oliet and Mothet, 2009). Recently, it has emerged that only D-SER gates NMDA receptors at the synaptic level, whereas extra-synaptic receptors are gated by glycine (Papouin *et al*., 2012; Sullivan and Miller, 2012). Furthermore, D-SER gates NMDA receptors only in the mature brain, whereas glycine gates NMDA receptor activity in the immature brain (Ferreira *et al*., 2017). In our study, we used adult rats with a weight of about 300-350 g. For these reasons, we chose D-SER as the more effectively co-agonist of NMDA receptors, with an expectation of a reduction in place field remapping. Accordingly, we found D-SER strongly reduced remapping (Day 2 vs Day 1). Moreover, after D-SER administration, we observed an increase in neuronal excitability in darkness. Indeed, NMDA receptor activation increased firing rates in claustral place cells when the rats navigated in darkness. We conclude that D-SER affects place fields in darkness, perhaps by increasing claustral place cell excitability.

Next, to assess the role of NMDA receptor co-activation in the local circuit, we investigated claustral oscillations following drug administration. Measurement of theta power (oscillations active during exploratory behaviour) revealed place field remapping in darkness was not driven by changes in the local network in the claustrum. Theta waves, which are present during various types of locomotor activities, have been assumed to be generated by the activation of NMDA receptors. Specifically, a combination of NMDA receptor blockers abolished all theta activity in the hippocampus (Buzsaki, 2002). Here, we observed theta power was not modulated by NMDA receptor activation in the claustrum.

The constant angle formed in darkness by claustral place cells and its substantial reduction following D-SER administration, led us to hypothesize that synaptic plasticity might mask an underestimated input to claustrum. Claustral place cell responses may have visual cortical inputs; however, transitioning from light to dark did not result in changes in activity in secondary visual cortex, but did reduce delta-band claustral oscillations. Delta oscillations may support attentional dynamics (Harmony, 2013); perhaps claustral remapping involves processes related to attention, rather than direct visual inputs. Here, we observed the presence of the distal beacon in the environment induces changes in the oscillatory activity of the visual cortex as well as the claustrum. Both brain regions were modulated in the delta-band by the distal beacon. We also observed that interactions between secondary visual cortex–claustral crosstalk in the delta and theta-band was not affected by the presence of the distal beacon.

Overall, our findings show that NMDA receptor activation stabilizes claustral place fields in darkness, perhaps by increasing the strength of synaptic transmission. Our findings of differential visual cortical and claustral activity during the light/dark protocol in the presence and absence of a distal beacon, expand on previous findings suggesting that claustral place cell remapping might be due to attention to distal beacon rather than direct inputs from the visual system. Future research might profitably focus on this issue to confirm the role of distal beacons on claustral cell remapping. (742 words)

## Material and Methods

### Subjects

A total of 15 male Lister-hooded rats (Envigo, United Kingdom) weighing 300-350 g were used. Upon arrival, they were housed two per cage and kept in a temperature-controlled laminar airflow unit with a 12/12 h light/dark cycle beginning at 8:00 A.M. with *ad libitum* access to food and water. They were handled daily for 10 days before electrode implantation. Rats were observed and handled during a recovery period of 10 days following electrode implantation. Before starting with electrophysiological recordings animals underwent a food deprivation procedure until they reached 85% of free-feeding body weight and were maintained at this weight during the entire study. In 3 of the animals a preliminary screening was performed using a single recording of 10 minutes where rats navigated in a square arena to ensure the presence of place cells in the claustrum. In 10 animals we performed electrophysiological recordings following microdrive (Axona, Ltd, UK) implantation in the claustrum using a light/dark/light protocol. In 6 of the 10 animals D-SER was systemically administrated. Additionally, 3 animals were used for LFP recordings to determine how effectively the 10mg/Kg dose D-SER influences the local claustral network. Finally, in 2 animals we performed electrophysiological recordings following microdrive implantation in both the claustrum and secondary visual cortex.

### Compliance with ethical standards

Experiments were conducted in accordance with locally-applicable laws and regulations, and international guidelines of good practice. Surgeries were conducted under isoflurane anaesthesia, an appropriate post-surgery monitoring and analgesia regime was in place, and every effort was done to minimize suffering.

### Behavioural testing

#### Place cells recordings in light/dark

The experimenter entered the recording room at the start and the end of each sequence of session to clean rat urine traces in the arena, for sugar pellet throwing and to disentangle the recording cable. Electrophysiological recordings were performed from rats navigating an 80×80 cm square and black arena (see Fig. 1a). The arena was in the middle of an opaque, black-curtained, square enclosure. The curtains were open/closed during recordings in light/dark respectively. All the equipment and devices (including the recording setup and computers) of the room were covered with black bags to reduce visual cues. During the recording in light the apparatus was indirectly lit by four symmetrically-positioned spotlights. During foraging a small amount of pellet (TestDiet^™^, 5 TUL, USA) was thrown in the arena at random locations.

### LFP spectra recording

After a recovery period from tetrode implantation in the claustrum, the head-stage was connected to the microdrive for recordings. Three rats were subcutaneously injected with normal saline (Day 1, NaCl, 0.9% 10 mg/Kg) and D-SER (Day 2, 10 mg/Kg, Sigma-Aldrich Ireland) and placed in a box (24h x 38w x 38l cm). The LFP recording started thirty minutes following the substance administration (Fig.5a). LFPs were recorded for thirty minutes during which sugar pellets were thrown in the box every five minutes. The experimenter was outside the room for the entire recording, except every five minutes for pellet throwing. The oscillatory activity from the secondary visual cortex and claustrum was simultaneously recorded during light/dark animal navigation without (Day 1) or with (Day 2) a distal visual beacon (flashing green light) at a distance of 2 meters (Fig. 6a) while animals navigated in a square arena.

### Electrode implantation

Following the acclimation period, animals underwent surgery for electrode implantation 25 µm above the claustrum to avoid tissue damages. Rats were implanted with bundles of eight tetrodes of ø 25 µm platinum-iridium wires with impedance 150-350 KΩ (California Fine Wire Ltd., CA, USA). Microdrives were built using eight tetrodes and implanted after craniotomy in the caudal part of the anterior claustrum, according to the atlas coordinates (Paxinos and Watson, 5^th^ Edition). The claustral coordinates used relative to bregma were: anterior-posterior (AP) + 2.52 mm, medial-lateral (ML) ± 2.00 mm, dorso-ventral (DV) -4.6 mm from top of the cortex, at an angle of 13^0^. Coordinates for stereotaxic implantation of a bipolar electrode (Stainless steel wire, SS-5T 127 µm, Science Products GmbH, Hofheim, DE) in secondary visual cortex relative to bregma were: anterior-posterior (AP) - 5.28 mm, medial-lateral (ML) ± 4.00 mm, dorso-ventral (DV) -1.4 mm from top of the cortex, at an angle of 24.8_0_. The bipolar electrode was connected to two microdrive electrodes.

### Screening and Recording

After the recovery period, tetrodes were lowered about 25 µm to the upper layer of the claustrum. A delay of 24 h was interposed between this stage and the screening sessions. During screening sessions, a cable connecting the recording system to the head-stage (Axona Ltd., UK) was plugged into the microdrive (Axona Ltd., UK). During rat navigation, a screening/recording was performed by using Axona system and software DacqUSB (Axona Ltd., UK). Tetrodes were lowered slowly through the brain (25 µm/day) until single unit signals were considered of sufficient amplitude, with spike properties typical of place cells (Table 1) and displaying a place field. Signals were amplified between 3000 and 12000 and bandpass filtered between 380 Hz and 7 kHz for single-unit detection. A camera was located above the arena allowing tracking of the animal’s head position. The animal’s path was visualized and recorded by the software DacqUSB (Axona Ltd., UK).

### Data analysis

#### Spike units

Graphical cluster-cutting Tint software (Axona Ltd., UK) was used to cluster-cut the waveforms selected. Place cells were considered as those cells with specific electrophysiological properties (ref., Table 1) that visually and statistically showed a place field. Based on their activity, unit identification involved further criteria. A percentage (16%) of the recorded place cells disappeared after the first session (L1), possibly due to positional instability of the electrodes and/or the cell. In this case, the cell was counted as a place cell according to its properties but without the following D/L2 sessions. Statistical analysEs for mean spike amplitude, mean spike-width, mean firing frequency, mean inter-spike interval (ISI), spatial information content (Skaggs) and spatial coherence (mean ± SEM) was performed (Table 1). The spike amplitude (µV) was measured by the difference between the first negative peak and first positive peak, spike-width (µs) was taken at 25% of the spike amplitude; firing frequency was considered as the number of spikes per second (Hz), the ISI (ms) was represented by the time between subsequent action potentials and the spatial information content was the information content of the spatial firing map. Spatial coherence was used as a measure of the place field remapping and represented the correlation between the raw firing map and smoothed firing map (value at each pixel is replaced the 8 neighbouring pixels of the non-smooth map). The measure of the total burst as the total number of burst events in a cell was used for bursting cells.

#### Local field potential

LFP traces obtained from claustrum were referenced to the animal ground and power was calculated using a random channel for each tetrode. Relative theta power (theta/total) was calculated for each session (L1, D and L2) by calculating Welch’s periodogram for the LFP signal and then integrating bands of the periodogram (5 - 11 Hz for theta) using Simpson’s method. Total power was calculated by integrating the periodogram between 1.5 - 90 Hz. This range was chosen to eliminate very low and very high frequency noise present in the LFP recording. Relative powers in these studies were calculated by dividing the power in a band giving a measure of how much of the overall LFP activity is accounted for by this frequency band. The range of relative powers was 1.5-4 Hz for delta, 5-11 Hz for theta, 30-90 Hz for gamma oscillations. Coherence was calculated to check if there was a consistent phase relationship between LFP oscillations in two different brain regions at certain frequencies. Coherence for two signals x and y, C_xy_ (f) is a function of frequency which returns a value between 0 and 1. A value of 1 indicates a constant phase relationship between the signals at the frequency f, while a value of 0 indicates no consistent phase relationship between the two signals at f. Coherence in a frequency band (e.g. delta 1.5-4 Hz) was estimated as the average of the sample coherence values in that band. We calculated the magnitude squared coherence, using Scipy, so C_xy_ (f) = abs (P_xy_ (f))^2^ / (P_xx_ (f) * P_yy_ (f)) where P_xx_ and P_yy_ are the power spectral density estimates of x and y, and P_xy_ is the cross spectral density estimate of x and y.

#### Spatial firing stability

To assess whether place cells had a stable place field between successive sessions, we computed, using NeuroChat (Islam *et al*., 2019) spatial coherence along the L1/D/L2 sequence to establish place field stability. In addition, the equation L1DL2 (angle) was used to assess the measure of the angle (degrees) between the place field in darkness (D) and the two place fields in light (L1 and L2) using the COM of each place field:

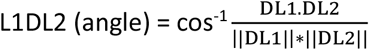

where DL1 and DL2 (numerators) are the vectors from the place field centroid in D to the place field centroid in L1 and from the place field centroid in D to the place field centroid in L2, respectively. DL1.DL2 is the dot product of the vectors and it is divided by the product between the norm of the vectors DL1 and DL2. The equation below was used to measure the difference in terms of distance (cm) between the place field centroids (x and y coordinates) in L1 and D, L1 and L2 and D and L2. The difference (cm) allowed to measure the shift in the place field between light and dark in control and following drug injection.

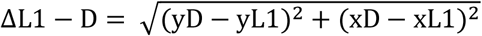

where x and y are the coordinates of the place field centroid, and the equation represents the distance formula.

#### Position and Speed estimation

DacqTrack (Axona, Ltd, UK) includes various methods for tracking the x and y coordinates of a moving target within the image. During L1/D/L2 and LFP recordings, two-spot (big/small) tracking method was used to track the rat’s position. The average running speed of the rat during navigation was measured in cm/sec.

### Histological analyses

Coronal brain sections were made in the coronal plane with a freezing microtome (50 µm sections in 4 series). One series was directly mounted on gelatin-subbed slides while the other was stored in a cryoprotective solution. The mounted sections were dried overnight, followed by rehydration in a series of ethanol solutions of decreasing concentrations (2 × 100%, 90%, 70%). The sections were then stained with cresyl violet after two minutes in deionized water. Following cresyl violet staining, sections were placed in deionized water and dehydrated in a series of ethanol solutions (70%, 90%, 2 x 100% series). Finally, mounted sections were defatted in xylene and coverslipped with DPX (ThermoFisher, Waltham, MA). The sections were imaged with Leica DM5000B microscope with a Leica DFC310FX digital camera and Leica Application Suite image acquisition software. Tetrode and bipolar electrode tracks were identified in the acquired images.

### Drugs

Rats were subcutaneously injected with NMDA receptor co-agonist D-SER (s.c. D-SER 10mg/Kg, Sigma-Aldrich, Ireland) in vehicle. D-SER was mixed in 0.9% of normal saline (NaCl) to achieve 10 mg/ml solution. Control rats received no injections. For LFP recordings, rats were injected with normal saline (NaCl, 0.9% w/x Sodium Chloride BP, Braun, Dublin) as a control for the following day when D-SER was administered.

### Experimental Design and Statistical Analysis

The experimental designs are illustrated in Figs.1a, 2a, 5a and 6a. The present study is divided in four main sections. Firstly, we investigated place cell activity when rats navigated in a square arena 2ith manipulation of visual stimuli. In this case, the experimental protocol comprised three sessions of 10 minutes each where rats (n rats = 10, n place cells = 19) moved from light (L1) to dark (D) and back to light (L2). Secondly, the protocol was repeated the following day (Day 2) at the same time as the previous day (Day 1). Day 1 was considered as the control for Day 2. The Day 2, a subgroup of rats (n rats = 6, n place cells = 9) received (subcutaneously) 10mg/Kg of NMDA receptor co-agonist D-SER thirty minutes before the recording session (see “Drug” section for further details). Thirdly, an additional group of rats (n = 3) were used to test the total power in the LFP and compared to saline. In this case, rats were subcutaneously injected with saline (Day 1) and D-SER (Day 2). Following the injection, rats were placed into a box where thirty minutes of LFP recording was performed (see “LFP spectra recordings” section for more details). For the larger part of the study, when the normal distribution was present, we ran parametric tests (t tests, paired), whereas nonparametric tests (Wilcoxon signed-rank test for paired data) were used otherwise. The Wilcoxon signed-rank test for paired data was used to compare theta power over the two days of recordings before and after the drug. The software GraphPad Prism 8 was used for statistical analysis (t test and Wilcoxon signed-rank test) and graphs. NeuroChaT (Islam et al., 2019) was used to analyse spike properties and relative theta power (see “spike units” and “LFP” sections, respectively). The analysis for the total power in the 30 minutes LFP recording was written in Python (v3.7), extending NeuroChaT. Finally, we investigated the influence of a distal visual beacon on the oscillatory activity of secondary visual cortex and claustrum. To this end, 2 implanted rats (n samples = 4, 2 LFP traces from the visual cortex (ctrl and V) and 2 from claustrum (ctrl and VC) were averaged for each rat) navigated in a square arena in light and dark (L1/D) in the absence (Day 1) or presence (Day 2) of a distal beacon in the environment. The analysis for the total power, the relative delta, theta, and gamma powers and coherency between visual cortex and claustrum was written in Python (v3.7).

## Notes

**Funding**: This work was supported by Science Foundation Ireland grant SFI 13/IA/2014.

### Competing Interest Statement

The authors have declared no competing interest.

